# A caspase-1-cathepsin AND-gate probe for selective imaging of inflammasome activation

**DOI:** 10.1101/2025.05.27.656501

**Authors:** Shiyu Chen, Una Goncin, Jiyun Zhu, Shih-Po Su, Traci Ann Czyzyk, Corin O. Miller, Raana Kashfi Sadabad, Matthew Bogyo

**Author notes:** Shiyu Chen and Una Goncin contributed equally.

## Abstract

Caspase-1 is a key mediator of the inflammasome pathway, which is associated with several inflammatory disorders including obesity, diabetes mellitus (DM), cardiovascular diseases (CVDs), cancers and chronic respiratory diseases. Although substrate-based probes can be used to visualize the activity of caspase-1, none are selective enough for use as imaging agents. Here, we report the design and synthesis of a AND-gate substrate probe (**Cas1-Cat-Cy7**) that requires processing by both caspase-1 and cathepsins to produce a signal. Because both enzymes are only found together and active at the site of inflammasome activation, the resulting probe can be used to image caspase-1 mediated inflammation. We demonstrate that the probe produces selective signals in *ex vivo* biochemical and cellular assays and in a mouse model of acute inflammation.

---

The inflammasome is a multiprotein complex crucial for innate immune responses^1^. The complex can be activated by pathogen-associated molecular patterns (PAMPs) or damage-associated molecular patterns (DAMPs). Upon activation, the inflammasome recruits and activates the protease caspase-1 through adapter proteins such as ASC (Apoptosis-associated speck-like protein containing a caspase activation and recruitment domain)^2^. Active caspase-1 cleaves gasdermin D, leading to the formation of pores in the cellular membrane and eventually a lytic form of cell death known as pyroptosis^3^. While appropriate inflammasome activation is critical for host defense, dysregulated inflammasome activity is linked to autoimmune diseases^4^, metabolic disorders^5^, cancers^6^, cardiovascular diseases^7^ and neurodegenerative conditions^8^. Hence, direct optical imaging of inflammasome activity has high value for understanding disease mechanisms and facilitating therapeutic development. Current optical probes for inflammasome activity rely only on caspase-1 activity^9-12^. These probes typically use a peptide sequence (e.g., YVAD or WEHD) linked to a fluorophore and quencher. Caspase-1 cleaves the substrate sequence, separating the quencher from the fluorophore, giving fluorescent signal (Figure 1a). Although substrate-based probes can be used to visualize the activity of caspase-1, selectivity is often limited by other enzymes that process the substrate. A cathepsin-caspase-1 AND-gate probe is a next-generation molecular tool designed to address the limitations of traditional caspase-1 probes by requiring activation from multiple enzymes that are only present and active together at sites of inflammasome mediated inflammation to generate a signal (Figure 1b). Cathepsins are a family of lysosomal proteases critical to cellular homeostasis and various physiological processes^13^. Notably, highly concentrated cathepsin activity is associated with sites of inflammation^14^. Active caspase-1 cleaves gasdermin D, resulting in pyroptosis. Pyroptic cells release IL-1β and IL-18^15^, which is important for the activation of macrophages^16^. These proinflammatory macrophages up-regulate the expression of cathepsins. In the simplest scheme, a **Cas1-Cat-Cy7** AND gate probe is first cleaved by active caspase-1 during pyroptosis, then is internalized and cleaved by lysosomal cathepsins produced by macrophages recruited to the inflammation site (Figure 1c).

**Figure 1.**
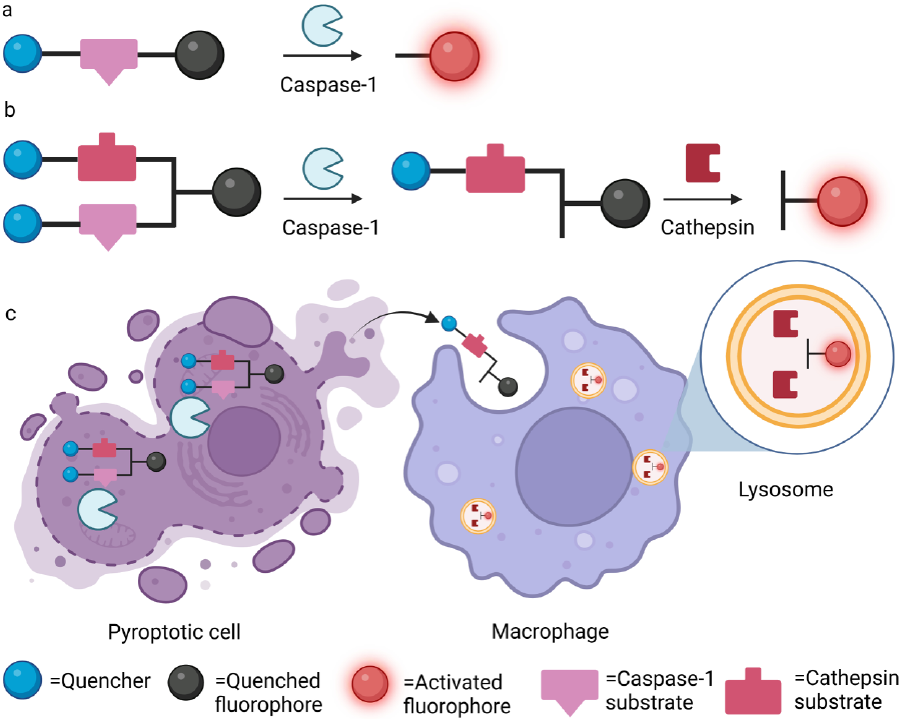
a). Scheme of single-substrate caspase-1 probe. b). Scheme of cathepsin-caspase-1 AND-gate probe. c). Cartoon of **Cas1-Cat-Cy7** AND-gate probe being cleaved by active caspase-1 and then internalized and cleaved by lysosomal cathepsins in macrophages.

An AND-gate probe targeting apoptotic caspases and cathepsins for imaging tumors has been described previously by Widen et al^17^. This probe utilizes a D-glutamic acid (D-Glu) as the central linker due to its high *in vivo* stability compared to L-glutamic acid (L-Glu)^17^. To generate a caspase-1/cathepsins AND-gate substrate, we attached the two substrate peptides containing the same quencher group to the two carboxylic groups of the D-glutamic acid linker. We then attached a fluorescent dye to the free amino group on the linker (Figure 2a).

**Figure 2.**
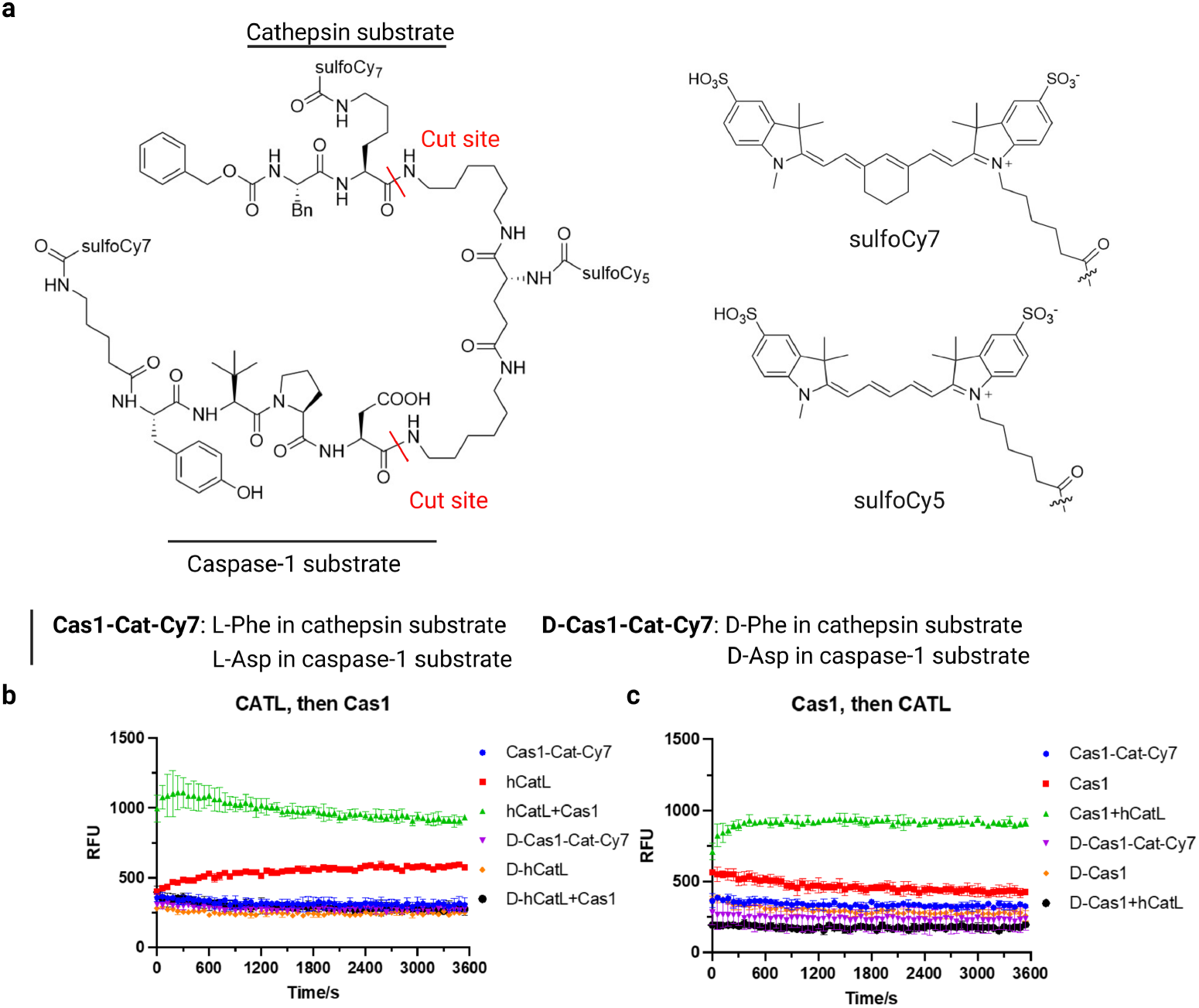
a). Structure of cathepsin-caspase-1 AND-gate probe **Cas1-Cat-Cy7** and the corresponding negative control **D-Cas1-Cat-Cy7. D-Cas1-Cat-Cy7** contains a D-Phe in the P2 position of the cathepsin sequence and a D-Asp in the P1 position of the caspase-1 sequence to block proteolytic cleavage by the respective protease. b). Progress curve of **Cas1-Cat-Cy7** incubated with cathepsin L followed by caspase-1 compared with negative control probe **D-Cas1-Cat-Cy.** c). Progress curve of **Cas1-Cat-Cy7** incubated with caspase-1 followed by cathepsin L compared with negative control probe **D-Cas1-Cat-Cy. Cas1-Cat-Cy7** and **D-Cas1-Cat-Cy** were used at 5 μM. Cathepsin L was used at 5 nM. Caspase-1 was used at 10 nM. RFU, relative fluorescence units. Data represents three replicates.

Fluorescence activation occurs only after both peptides have been cleaved by caspase-1 and cathepsins. We chose to use a non-natural peptide sequence that has previously been shown to be specifically processed by caspase-1 (Asp-Pro-tertLeu-Tyr)^18^ and the previously validated peptide substrate that is cleaved efficiently by cysteine cathepsins (Lys-Phe)^19^. Although these substrates are quite selective for their intended proteases, application of single-substrate probes *in vivo* is limited by background non-specific activation. The AND-gate strategy reduces this non-specific activation by requiring both enzymes to be activated at the site of inflammation. We chose to use a sulfoCy5-sulfoCy7 fluorophore-quencher pair due to the potential use for ratiometric imaging^20^. Consistent with our previous design, the final fluorescent fragment produced after protease cleavage contains two free amine groups that remain protonated and retained in low pH lysosomes of macrophages.

We synthesized the caspase-1-cathepsin AND-gate probe, **Cas1-Cat-Cy7**. We also synthesized a negative control probe **D-Cas1-Cat-Cy7**, in which the natural L-Asp in Caspase-1 substrate P1 is replaced by D-Asp and L-Phe in the Cathepsin substrate P2 is replaced by D-Phe to prevent cleavage by either protease. To test the probes, we incubated them with recombinant human cathepsin L (hCatL) and caspase-1 either separately or sequentially in combinations (Figure 2b-c). Notably, we found that the **Cas1-Cat-Cy7** probe produced an enhanced fluorescence signal only when both cathepsin and caspase-1 were incubated sequentially. As expected, we found negative control probe **D-Cas1-Cat-Cy7** did not show any fluorescence increase after incubation with the two enzymes.

We next evaluated the **Cas1-Cat-Cy7** and **D-Cas1-Cat-Cy7** in a co-culture of Ea.hy926 and RAW 264.7 cells. Ea.hy926 were incubated with Val-boroPro (50 μM) for 2h to induce pyroptosis. It has been shown previously that RAW 264.7 cells under VBP stimulation do not express active caspase-1^21^. **Cas1-Cat-Cy7** or **D-Cas1-Cat-Cy7** were added to the cells for 2h before acquiring images. We observed significant fluorescence signal within 2h. As expected, **Cas1-Cat-Cy7** was processed and produced a fluorescent signal in the RAW 264.7 macrophage cells only when VBP was added and both cells were present, while **D-Cas1-Cat-Cy7** did not produce a fluorescent signal in any conditions (Figure 3a and Figure S1, S2). Furthermore, EA.hy926 cells treated with **Cas1-Cat-Cy7** in the absence of RAW cells (Figure 3c) or VBP (Figure 3b) did not show any fluorescence signals. Finally, neither probe showed any activation with EA.hy926 cells in which caspase-1 was knocked out (Figures S3, S4). These results suggested that the **Cas1Cat-Cy7** probe is cleaved by caspase-1 expressed by Ea.hy926 cells during pyroptosis, and then is internalized and cleaved by lysosomal cathepsins in the macrophages, where the fluorescent signal is retained in the lysosomes.

**Figure 3.**
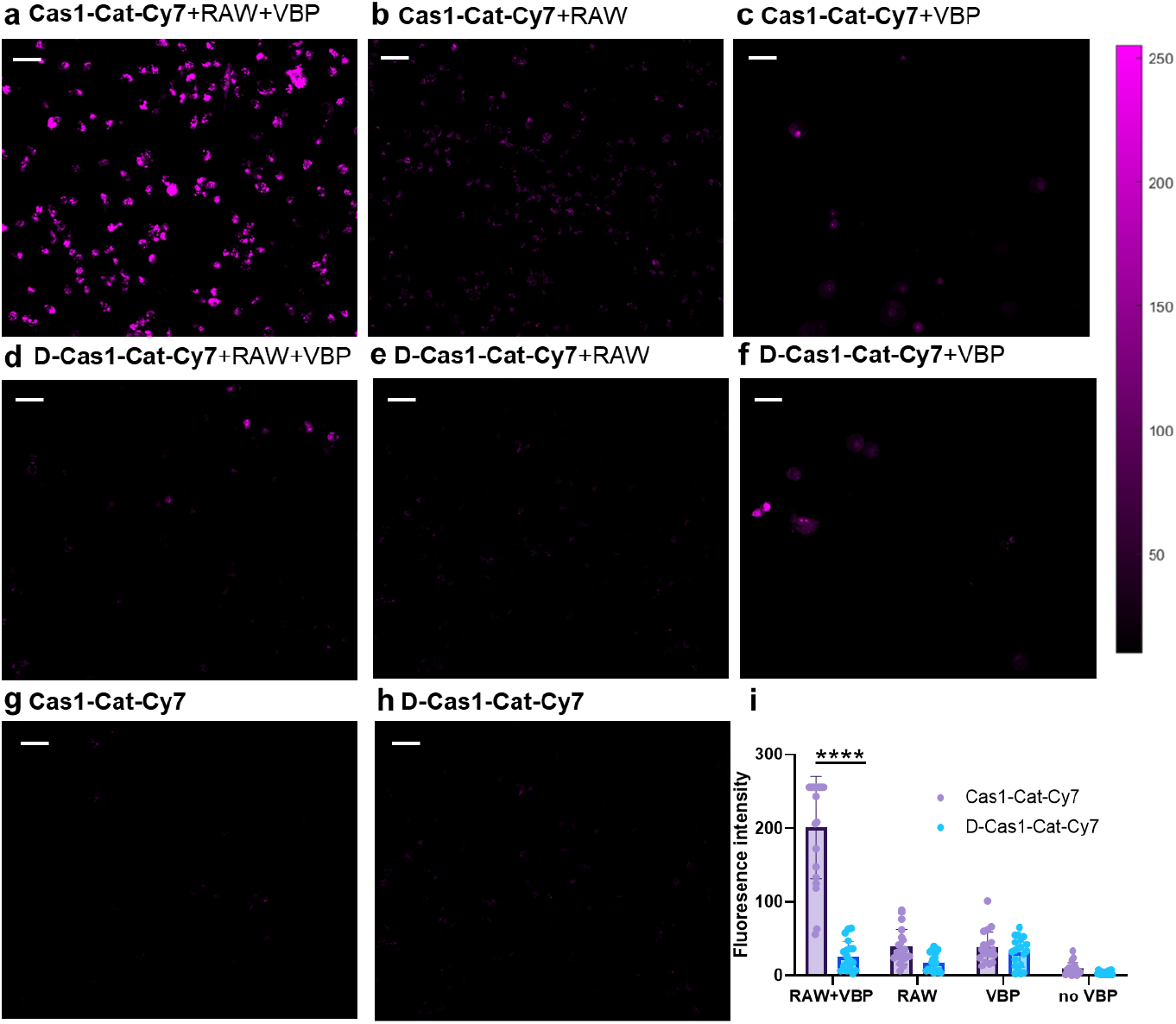
Representative fluorescence microscopy images of EA.hy926 cells cocultured with RAW 264.7 cells labelled with either **Cas1-Cat-Cy7** or **D-Cas1-Cat-Cy7** under Leica MICA Microhub microscope (20x). Cells were incubated with Val-boroPro (50 μM) for 2h before probe (1 μM) was added and incubated for another 2h. Fluorescence was collected at Cy5 channel. Scale bar: 20 μm. a). Coculture of EA.hy926 and RAW 264.7 cells were incubated with Val-boroPro (50 μM) for 2h and then **Cas1-Cat-Cy7** (1 μM) was added and incubated for another 2h. b). Coculture of EA.hy926 and RAW 264.7 cells were incubated with **Cas1-Cat-Cy7** (1 μM) for 2h. c). EA.hy926 cells were incubated with Val-boroPro (50 μM) for 2h and then **Cas1-Cat-Cy7** (1 μM) was added and incubated for another 2h. d). Coculture of EA.hy926 and RAW 264.7 cells were incubated with Val-boroPro (50 μM) for 2h and then **D-Cas1-Cat-Cy7** (1 μM) was added and incubated for another 2h. e). Coculture of EA.hy926 and RAW 264.7 cells were incubated with **D-Cas1-Cat-Cy7** (1 μM) for 2h. f). EA.hy926 cells were incubated with Val-boroPro (50 μM) for 2h and then **D-Cas1-Cat-Cy7** (1 μM) was added and incubated for another 2h. g). EA.hy926 cells were incubated with **Cas1-Cat-Cy7** for 2h. h). EA.hy926 cells were incubated with **D-Cas1-Cat-Cy7** for 2h. i). Quantification of the fluorescence signal from microscopy images. N=20. Statistical analysis was performed using two-way analysis of variance (ANOVA), ****P < 0.0001.

We next evaluated the **Cas1-Cat-Cy7** and **D-Cas1-Cat-Cy7** probes in an LPS-ATP induced mouse model of inflammation. Sixteen female B6NTac mice were purchased from Taconic (Germantown, NY, USA) and housed in a 12:12 light-dark cycle with standard rodent chow and water provided *ad libitum*. Mice (n=10) were anesthetized (using 2% isoflurane) and acute inflammation was induced using a subcutaneous injection of LPS and ATP (LPS+ATP; 5 ug and 5 mg, respectively, in 30 μL deionized water) in the dorsal surface of the left hind paw. A subset of mice which not treated with LPS+ATP (n=6) served as controls^10^. Maximum fluorescent activation was observed at 8h.

As shown in Figure 4, mice treated with the **Cas1-Cat-Cy7** probe showed significantly increased fluorescence signal over baseline in the LPS/ATP treated paw (left paw) compared to the untreated paw (right paw). In addition, the signal in mice treated with the **Cas1-Cat-Cy7 probe** was significantly increased compared to mice treated with the negative control probe **D-Cas1-Cat-Cy7**, suggesting that signal enhancement was due to protease activity. However, we did note that the negative control probe had relatively high background signal in the LPS/ATP treated paws. It is likely that this background signal is the result of accumulation of the un-cleaved probe caused by increased blood flow at the site of inflammation. Ratiometric imaging offers potential to reduce the impact of such background signals because the probe can be calibrated using a ratio of the fluorescent intensities of two emission bands (cy5 and cy7)^20^. This eliminates background signals from the un-cleaved probe that might otherwise be interpreted as enzyme activity.

**Figure 4.**
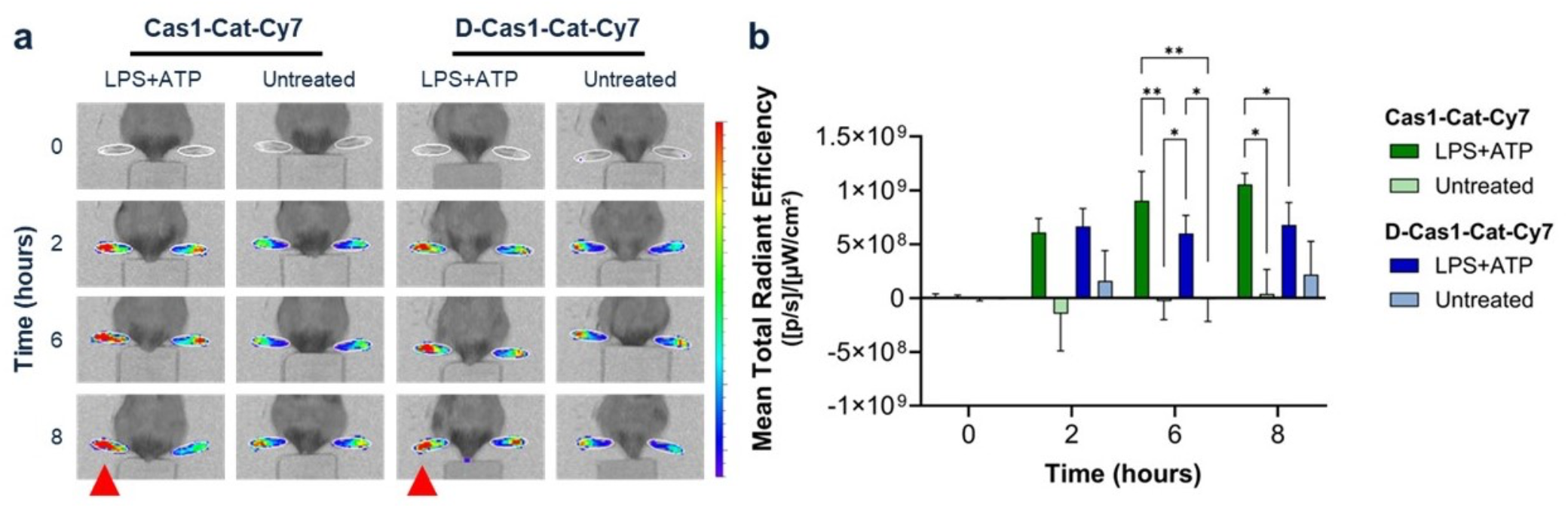
*In vivo* optical imaging of acute hindpaw inflammation using **Cas1-Cat-Cy7**. a). Representative *in vivo* fluorescence images of mice that were untreated (n=6) or injected with LPS+ATP (n=10, 5 ug and 5 mg, respectively) in the dorsal surface of the left hindpaw (red arrowhead). **Cas1-Cat-Cy7** and **D-Cas1-Cat-Cy7** probes were administered intravenously, and images were collected over an 8-hour period using a Cy5 filter (*ex*=649 nm, *em*=666 nm) on an IVIS SpectrumCT. b). Comparison of the mean total radiant efficiency ([p/s]/[μW/cm^2^; calculated based on signal difference between the left and right hindpaw of each mouse) using the **Cas1-Cat-Cy7** and **D-Cas1-Cat-Cy7** in LPS+ATP treated and untreated mice. A two-way analysis of variance (ANOVA), followed by Tukey post-hoc multiple comparison test was used to compare differences in total radiant efficiency between LPS+ATP treated and untreated mice using **Cas1-Cat-Cy7** and **D-Cas1-Cat-Cy7** over time.

In summary, we describe the synthesis, characterization, and biological application of a **Cas1-Cat-Cy7** AND-gate probe for selective imaging of inflammasome activity *in vivo*. We demonstrate in a EA.hy926 cell pyroptosis model as well as LPS-ATP paw inflammation mouse model that the probe shows selective activation that is dependent on both caspase-1 and cathepsin activity. Our data suggests that **Cas1-Cat-Cy7** can be used for selective *in vivo* imaging of inflammasome activity. We believe this will be particularly valuable for pre-clinical testing of therapeutics designed to treat inflammation-related diseases.

## Data Availability

Data that support the findings of this study are available in the supplementary materials of this article.

## Acknowledgements

The authors thank Salomon Martinez and Mahamudul Haque for technical assistance in performing the mouse imaging studies. The authors thank Kasia Groborz for valuable inputs in cell experiment design. This work was supported in part by NIH grant (R01 EB028628 to M.B.).

## AUTHOR INFORMATION

### Author Contributions

The manuscript was written through contributions of all authors.

---

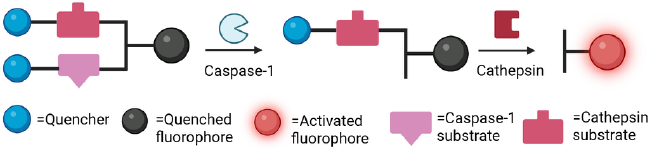

## REFERENCES

1. D. Zheng, T. Liwinski, E. Elinav. Inflammasome activation and regulation: toward a better understanding of complex mechanisms. Cell. Discov. 2020, 6, 36.

2. N. B. Bryan, A. Dorfleutner, Y. Rojanasakul, C. Stehlik, J. Activation of inflammasomes requires intracellular redistribution of the apoptotic speck-like protein containing a caspase recruitment domain. Immunol. 2009, 182, 3173–3182.

3. E. A. Miao, J. V. Rajan, A. Aderem. Caspase-1-induced pyroptotic cell death. Immunol. Rev. 2011, 243, 206–214.

4. D. Pohl, S. Benseler. Systemic inflammatory and autoimmune disorders. Handb Clin. Neurol. 2013, 112, 1243–1252.

5. K. Esposito, D. Giugliano. The metabolic syndrome and inflammation: association or causation? Nutr. Metab. Cardiovasc. Dis. 2004, 14, 228–232.

6. H. Zhao, L. Wu, G. Yan, Y. Chen, M. Zhou, Y. Wu, Y. Li. Inflammation and tumor progression: signaling pathways and targeted intervention. Signal. Transduct. Target. Ther. 2021, 6, 263.

7. M. Y. Henein, S. Vancheri, G. Longo, F. Vancheri. The role of inflammation in cardiovascular disease. Int. J. Mol. Sci. 2022, 23.

8. W. Zhang, D. Xiao, Q. Mao, H. Xia. Role of neuroinflammation in neurodegeneration development. Signal. Transduct. Target. Ther. 2023, 8, 267.

9. X. Ren, M. Tao, X. Liu, L. Zhang, M. Li, Z. Hai. Caspase-1responsive fluorescence biosensors for monitoring endogenous inflammasome activation. Biosens. Bioelectron. 2023, 219, 114812.

10. Y. J. Ko, J. W. Lee, E. J. Yang, N. Jang, J. Park, Y. K. Jeon, J. W. Yu, N. H. Cho, H. S. Kim, I. Chan Kwon. Non-invasive in vivo imaging of caspase-1 activation enables rapid and spatiotemporal detection of acute and chronic inflammatory disorders. Biomaterials. 2020, 226, 119543.

11. H. Lin, H. Yang, S. Huang, F. Wang, D. M. Wang, B. Liu, Y. D. Tang, C. J. Zhang. Caspase-1 specific light-up probe with ag-gregation-induced emission characteristics for inhibitor screening of coumarin-originated natural products. ACS. Appl. Mater. Interfaces. 2018, 10, 12173–12180.

12. T. Liu, Y. Yamaguchi, Y. Shirasaki, K. Shikada, M. Yamagishi, K. Hoshino, T. Kaisho, K. Takemoto, T. Suzuki, E. Kuranaga, O. Ohara, M. Miura. Caspase-1 engagement and TLR-induced c-FLIP expression suppress ASC/caspase-8-dependent apoptosis by inflammasome sensors NLRP1b and NLRC4. Cell. Rep. 2014, 8, 974–982.

13. T. Yadati, T. Houben, A. Bitorina, R. Shiri-Sverdlov. The ins and outs of cathepsins: Physiological function and role in disease management. Cells 2020, 9.

14. S. Conus, H. U. Simon. Cathepsins: key modulators of cell death and inflammatory responses. Biochem. Pharmacol. 2008, 76, 1374–1382.

15. T. Bergsbaken, S. L. Fink, B. T. Cookson. Pyroptosis: host cell death and inflammation. Nat. Rev. Microbiol. 2009, 7, 99–109.

16. K. Nakanishi, T. Yoshimoto, H. Tsutsui, H. Okamura. Interleukin-18 regulates both Th1 and Th2 responses. Annu. Rev. Immunol. 2001, 19, 423–474.

17. J. C. Widen, M. Tholen, J. J. Yin, A. Antaris, K. M. Casey, S. Rogalla, A. Klaassen, J. Sorger, M. Bogyo. AND-gate contrast agents for enhanced fluorescence-guided surgery. Nat. Biomed. Eng. 2021, 5, 264–277.

18. A. W. Puri, P. Broz, A. Shen, D. M. Monack, M. Bogyo. Caspase-1 activity is required to bypass macrophage apoptosis upon Salmonella infection. Nat. Chem. Biol. 2012, 8, 745–747.

19. J. J. Yim, S. Harmsen, K. Flisikowski, T. Flisikowska, H. Namkoong, M. Garland, N. S. van den Berg, J. G. Vilches-Moure, A. Schnieke, D. Saur, S. Glasl, D. Gorpas, A. Habtezion, V. Ntziachristos, C. H. Contag, S. S. Gambhir, M. Bogyo, S. Rogalla. A protease-activated, near-infrared fluorescent probe for early endoscopic detection of premalignant gastrointestinal lesions. P. Natl. Acad. Sci. USA. 2021, 118.

20. F. F. Faucher, K. J. Liu, E. D. Cosco, J. C. Widen, J. Sorger, M. Guerra, M. Bogyo. Protease activated probes for real-time ratiometric imaging of solid tumors. Acs. Central. Sci. 2023, 9, 1059–1069.

21. M. C. Okondo, D. C. Johnson, R. Sridharan, E. B. Go, A. J. Chui, M. S. Wang, S. E. Poplawski, W. Wu, Y. Liu, J. H. Lai, D. G. Sanford, M. O. Arciprete, T. R. Golub, W. W. Bachovchin, D. A. Bachovchin. DPP8 and DPP9 inhibition induces pro-caspase-1-dependent monocyte and macrophage pyroptosis. Nat. Chem. Biol. 2017, 13, 46–53.

