## Supplementary material for "A caspase-1-cathepsin AND-gate probe for selective imaging of inflammasome activation": Supplmental Methods and figures

Table of Contents

1. General methods……………………………………………………………………...S2
2. Biology Methods……………………………………………………………………S2
3. Animal model………………………………………………………………………..S2
4. Synthesis of Cas1-Cat-Cy7 and D-Cas1-Cat-Cy7………………………………S3
5. Cell imaging of Ea.hy926 and Ea.hy926 KO cells………………………………S6
6. LCMS of compounds………………………………………………………………S8
7. References…………………………………………………………………………..S15

### 1. General methods

All commercial materials were purchased and used without further purification. SulfoCy5NHS and sulfoCy7NHS were purchased from Lumiprobe without further purification. Solvents with Sure-Seal bottles were purchased from Aldrich. The ESI mass spectra were obtained with an Agilent G6125B LCMS and Agilent 1100 LCMS using $H_2O$/ACN solutions. Dry solvents were obtained from a dry solvent system using alumina columns. Air or moisture sensitive reactions were conducted using Schlenk line technique. No unexpected or unusually high safety hazards were encountered.

### 2. Biology Methods

General cell culture

Ea.hy926 cells (ATCC CRL-2922) and Ea.hy926 caspase-1 KO cells (generous gift from Genentech) were cultured in Dulbecco's Modified Eagle's Medium (DMEM, Gibco, 11965-092) containing 4.5 g $l^{-1}$ of glucose, 0.3 g $ml^{-1}$ of L-glutamine, andsupplemented with 10% FBS and 100 U $ml^{-1}$ penicillin and 100 µg $ml^{-1}$ streptomycin. RAW 264.7 (ATCC TIB-71) macrophages were cultured in Dulbecco's modified Eagle's medium (DMEM, Gibco, 11965-092) containing 4.5 g $l^{-1}$ of glucose supplemented with 10% fetal bovine serum (FBS, GeminiBio, 100602), and 100 U $ml^{-1}$ penicillin and 100 µg ml−1 streptomycin (Gibco,15140-122). In general, Ea.hy926 cells and RAW246.7 were passaged 3 times after thawing before confocal microscopy.

Fluorogenic Substrate Cleavage Assay

Recombinant cysteine cathepsin L was from R&D Systems (Catalog No.: 952-CY). Recombinant caspase-1 was from Enzo Life Sciences. Buffers used for fluorogenic substrate cleavage assays were made as previously described.[1,2,3] All assays were conducted in clear Eppendorf tubes. Immediately before fluorescence measurements, protease was added to the final concentration reported in the text. Fluorescence was performed with a Cytation 3 imaging reader. Cy5 signal was detected by exciting at 640 nm and fluorescence emission was recorded at 690 nm. All experiments were repeated 3 times.

Procedures involving the care and use of animals in this study were reviewed and approved by the Institutional Animal Care and Use Committee at Merck & Co. During the study, the care and use of animals were conducted in accordance with the principles outlined in guidance from the Association for Assessment and Accreditation of Laboratory Animal Care, the Animal Welfare Act, the American Veterinary Medical Association Euthanasia Panel on Euthanasia, and the Institute for Laboratory Animal Research Guide to the Care and Use of Laboratory Animals.

### 3. Animal Model

All animal procedures at Merck (South San Francisco, CA, USA) were approved by the Institutional Animal Care and Use Committee (Protocol #4003344). Sixteen female B6NTac mice were purchased from Taconic (Germantown, NY, USA) and housed in a 12:12 light-dark cycle with standard rodent chow and water provided ad libitum. Mice (n=10) were anesthetized (using 2% isoflurane) and acute inflammation was induced using a subcutaneous injection of LPS and ATP (LPS+ATP; 5 µg and 5 mg, respectively, in 30 µL deionized water) in the dorsal surface of the left hindpaw. A subset of mice (n=6) served as controls, which were not treated with LPS+ATP. Due to the known decreased body temperature associated with LPS+ATP treatment, all mice were provided thermal support throughout the experiment and monitored closed.

Mice were placed on low autofluorescence diet (#AIN-76; Inotiv, Indianapolis, IN, USA) at least 4 days prior to optical imaging. **Cas1-Cat-Cy7** and **D-Cas1-Cat-Cy7** were provided in 10 mM stock solutions. For in vivo imaging, the probes were prepared as follows: 0.5 % 10 mM stock solution + 30 % PEG300 + 9.5 % DMSO + 60 % PBS. Each mouse received either **Cas1-Cat-Cy7** and **D-Cas1-Cat-Cy7** (5 nmol in 100 µL/mouse) intravenously via tail vein, approximately 15 minutes after LPS+ATP treatment. Mice were imaged using a Cy5 filter (ex = 649 nm, em = 666 nm) on an IVIS SpectrumCT (Caliper Life Science, Hopkinton, MA, USA) at 2 hours, 6 hours, and 8 hours post-probe injection. Settings (exposure, binning, f/stop, filters, lamp voltage, field of view) were kept constant throughout image acquisition. A region of interest (ROI) was drawn around the left and right hindpaw of each mouse using Living Imaging Software (Version 4.7.4; Revvity, Waltham, MA, USA). Normalized fluorescent intensity was expressed as the total radiant efficiency ([p/s]/[µW/cm²]) representing the signal difference between the left hindpaw compared to the right hindpaw. Results as expressed as mean total radiant efficiency of replicate mice ± SD.

At 8 hours post-probe injection, the total radiant efficiency of the **Cas1-Cat-Cy7** was significantly higher (1.5-fold; 1.05 x 109 ± 1.03 x 108) than that of the **D-Cas1-Cat-Cy7** (6.77 x $10^8$ ± 2.09 x $10^8$; p=0.043) in LPS+ATP treated mice. In the untreated mice, there were no significant differences in signal between probes over the 8-hour period. Comparing Probes: **Cas1-Cat-Cy7** signal was significantly higher in LPS+ATP treated mice compared to untreated mice at 6 hours (9.02 x $10^8$ ± 2.73 x $10^8$ and -2.97 x $10^7$ ± 1.71 x $10^8$, respectively; p=0.004) and 8 hours (1.05 x $10^9$ ± 1.03 x $10^8$ and 3.77 x $10^7$ ± 2.28 x $10^8$, respectively; p=0.027) post-probe injection. This increased signal was observed with the negative probe at 6 hours post-probe injection when comparing LPS+ATP treated mice (6.77 x $10^8$ ± 2.09 x $10^8$) to untreated mice (2.18 x $10^8$ ± 3.07 x $10^8$; p=0.045).

### Synthesis of Cas1-Cat-Cy7 and D-Cas1-Cat-Cy7

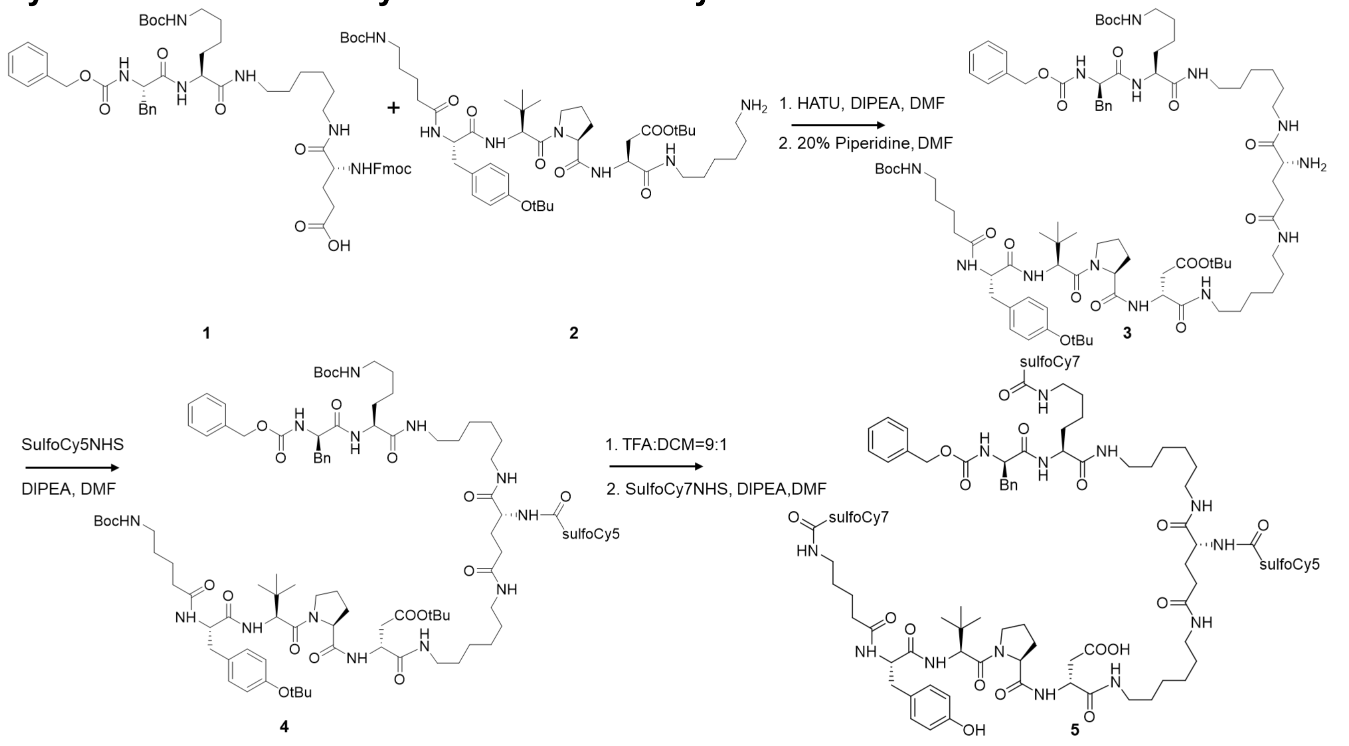

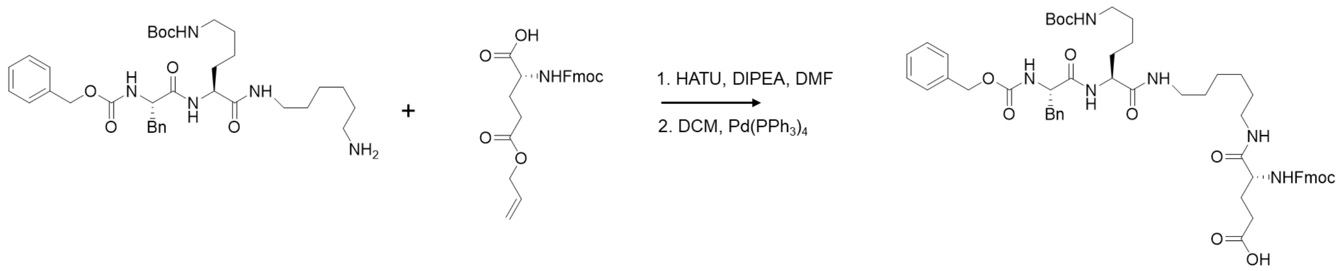


**Scheme S1.** Synthesis route for **Cas1-Cat-Cy7**.

HATU: Hexafluorophosphate azabenzotriazole tetramethyl uronium
DMF: Dimethylformamide
DCM: Dichloromethane
DIPEA: Diisoprophylethylamine

Synthesis of **1**
Intermediate **1** was synthesized according to the literature.[1,2] Briefly, 140 mg of **6**, 91 mg of **7** and 85 mg HATU, 96 μL of DIPEA were dissolved in 2 mL DMF. The reaction mixture was stirred for 2h. Pour the reaction mixture in ice water and the precipitate was collected by filtration. The precipite was redissolved in 5 mL DCM. 6.2 mg of $Pd(PPh_3)_4$ and 66 μL of Phenylsilane were added to the reaction mixture. Stirred the reaction for 1 hour. The solvent was then removed under vacuum and the residue was redissolved in 1 mL DMSO and purified with a reverse phase Combiflash with a gradient program of 5% MeCN:H2O (0.1% TFA) for 0-2 min, 5-90% for 2-23 min, 90-95% for 23-25 min. Purified fractions were collected and lyophilized to obtain a white powder (87 mg, 40% over two steps).

Synthesis of **2**
Compound **2** was synthesized by solid-phase synthesis according to the literature.[1,2]
512 mg of 2-Chlorotrityl Chloride Resin was swelled in DCM in a 60 mL solid phase extraction (SPE) tube for 20 min. DCM was then drained and washed with DCM two times. 796 mg of Fmoc-1,6-diaminohexane hydrochloride, 818 μL of DIEPA, 14 mL of DMF were mixed in a 20 mL vial and heated to dissolve. The dissolved solution was added to SPE tube, and the tube was shaken overnight. Solvent was then drained, and 10 mL of methanol was added to cap the unreacted resin. Wash with DMF three times, DCM three times, DMF three times and DCM three times. 10 mL of 20 % piperidine in DMF was added and shaken for 10 min. Drain the liquid, add another 10 mL 20 % piperidine in DMF and shake for 10 min. Rinse with DMF three times, DCM three times, and DMF three times. 1578 mg of Fmoc-Asp(Boc)-OH, 892 mg of HBTU, 816 μL of DIPEA were dissolved in 15 mL of DMF. Add the dissolved mixture to the SPE tube and shake for 1h. Drain the solvent, and rinse with DMF three times, DCM three times, and DMF for three times. 10 mL of 20 % piperidine in DMF was added and shaken for 10 min. Drain the liquid, add another 10 mL 20 % piperidine in DMF and shake for 10 min. Rinse with DMF three times, DCM three times, and DMF three times. 1298 mg of Fmoc-Pro-OH, 892 mg of HBTU, 816 μL of DIPEA were dissolved in 15 mL of DMF. Add the dissolved mixture to the SPE tube and shake for 1h. Drain the solvent, and rinse with DMF three times, DCM three times, DMF for three times and DCM three times. Repeat the Fmoc deprotection and rinse procedure. 1357 mg of Fmoc-tertLeu-OH, 892 mg of HBTU, 816 μL of DIPEA were dissolved in 15 mL of DMF. Add the dissolved mixture to the SPE tube and shake for 1h. Drain the solvent, and rinse with DMF three times, DCM three times, DMF for three times and DCM

three times. Repeat the Fmoc deprotection and rinse procedure. 1760 mg of Fmoc-Tyr(tBu)-OH, 892 mg of HBTU, 816 µL of DIPEA were dissolved in 15 mL of DMF. Add the dissolved mixture to the SPE tube and shake for 1h. Repeat the Fmoc deprotection and rinse procedure. 1760 mg of Fmoc-Tyr(tBu)-OH, 892 mg of HBTU, 816 µL of DIPEA were dissolved in 15 mL of DMF. Add the dissolved mixture to the SPE tube and shake for 1h. Drain the solvent, and rinse with DMF three times, DCM three times, DMF for three times and DCM three times. Repeat the Fmoc deprotection and rinse procedure. 833 mg Boc-5-Ava-OH, 892 mg of HBTU, 816 µL of DIPEA were dissolved in 15 mL of DMF. Add the dissolved mixture to the SPE tube and shake for 1h. Drain the solvent, and add 2 mL of Hexafluoroisopropanol (HFIP) in 6mL DCM to the SPE tube. Let the tube stay still for 2h. Drain the liquid and collect liquid in a 50 mL falcon tube. Air blow the solvent and add 5 mL of $H_2O$/CAN=1:1 solution to the residue. Freeze the solvent in liquid nitrogen and put the tube on lyophilizer to achieve 150 mg white power. The peptide was used without further purification.

Synthesis of **3**

The quantity 115 mg of **1** (0.117 mmol), 107 mg of **2** (0.117 mmol), 45 mg of HATU and 65 µL of DIPEA were mixed in a vial. 2 mL of anhydrous DMF was added to the mixture. The mixture was stirred for 2h and monitored by LC-MS. 0.2 mL of piperidine was added. The reaction mixture was stirred for 15 min and evaporated under vacuum. The crude was purified by a reverse phase Combiflash with a gradient of 10% MeCN: $H_2O$ (0.1% TFA) for 0-2 min, 10- 80% for 2-19 min, 80-95% for 19- 25 min. The purified fractions were collected and lyophilized to obtain a white powder (44 mg, 24% yield over two steps). HRMS calc. for $[C_{87}H_{138}N_{13}O_{18}]^+$ 1653.0277 found 1653.0251.

Synthesis of **4**
Intermediate **3** (44 mg, 26 µmol, 1 equiv.) and SulfoCy5NHS (19 mg, 26 µmol, 1.2 equiv.) were dissolved in DMF (500 µL). Then, DIPEA (44 µL, 0.26 mmol, 10 equiv.) was added and the reaction was agitated for 24 h at RT. The reaction mixture was evaporated under vacuum and redissolved in 1 mL of 1:1 MeCN: H2O (0.1% TFA). The crude was purified by a reverse phase Combiflash with a gradient program of 10% MeCN:H2O (0.1% TFA) for 0-2 min, 10-95% for 2-23 min, 95% for 23-26 min. The purified fractions were collected and lyophilized to give a blue powder (30 mg, 50% yield). HRMS calc. for $[C_{119}H_{173}N_{15}O_{25}S_2]^+$ 2278.2280 found 2278.2200.

Synthesis of **5** (**Cas1-Cat-Cy7**)
The quantity 15 mg of **4** (6.5 µmol) was dissolved in 1 mL of TFA solution (TFA: DCM = 9:1). The mixture was stirred at room temperature for 4 hours. The reaction mixture evaporated under vacuum. The crude was used without further purification.

The crude was combined with s SulfoCy7NHS (10.9 mg, 13 µmol, 2.0 equiv.) and then dissolved in DMSO (1000 µL). Then, DIPEA was added (11.2 µL, 65 µmol, 10 equiv.) and the reaction was agitated for 24 h at 37 °C. The reaction was evaporated and redissolved in 1 mL of 1:1 MeCN:$H_2O$ (0.1% TFA) and purified with a reverse phase Combiflash with a gradient program of 5% MeCN:$H_2O$ (0.1% TFA) for 0-2 min, 5-90% for 2-23 min, 90-95% for 23-25 min. Purified fractions were collected and lyophilized to obtain a blue powder (15 mg, 70% yield). HRMS calc. for $[C_{175}H_{228}N_{19}O_{35}S_6]^{3+}$ 1115.8318 found 2278.8317.

Negative control molecules **D-3**, **D-4** and **D-5** were synthesized following the same protocol, where L-Phe and L-Asp were substituted by D-Phe and D-Asp during solid phase peptide synthesis.

HRMS calc. for **D-5** $[C_{175}H_{228}N_{19}O_{35}S_6]^{3+}$ 1115.8318 found 1115.8318.

Synthesis of **6**
512 mg of 2-Chlorotrityl Chloride Resin was swelled in DCM in a 60 mL solid phase extraction (SPE) tube for 20 min. DCM was then drained and washed with DCM two times. 796 mg of Fmoc-1,6-diaminohexane hydrochloride, 818 µL of DIEPA, 14 mL of DMF were mixed in a 20 mL vial and heated to dissolve. The dissolved solution was added to SPE tube, and the tube was shaken overnight. Solvent was then drained, and 10 mL of methanol was added to cap the unreacted resin. Wash with DMF three times, DCM three times, DMF three times and DCM three times. 10 mL of 20 % piperidine in DMF was added and shaken for 10 min. Drain the liquid, add another 10 mL 20 % piperidine in DMF and shake for 10 min. Rinse with DMF three times, DCM three times, and DMF three times. 1102 mg of Fmoc-Lys(Boc)-OH, 892 mg of HBTU, 816 µL of DIPEA were dissolved in 15 mL of DMF. Add the dissolved mixture to the SPE tube and shake for 1h. Drain the solvent, and rinse with DMF three times, DCM three times, and DMF for three times. 10 mL of 20 % piperidine in DMF was added and shaken for 10 min. Drain the liquid, add another 10 mL 20 % piperidine in DMF and shake for 10 min. Rinse with DMF three times, DCM three times, and DMF three times. 704 mg of Z-Phe-OH, 892 mg of HBTU, 816 µL of DIPEA were dissolved in 15 mL of DMF. Add the dissolved mixture to the SPE tube and shake for 1h. Drain the solvent, and rinse with DMF three times, DCM three times, DMF for three times and DCM three times. Add 2 mL of Hexafluoroisopropanol (HFIP) in 6mL DCM to the SPE tube. Let the tube stay still for 2h. Drain the liquid and collect liquid in a 50 mL falcon tube. Air blow the solvent and add 5 mL of $H_2O$/CAN=1:1 solution to the residue. Freeze the solvent in liquid nitrogen and put the tube on lyophilizer to achieve 140 mg white power. The peptide was used without further purification.

### 4. Cell imaging of Ea.hy926 and Ea.hy926 KO cells.

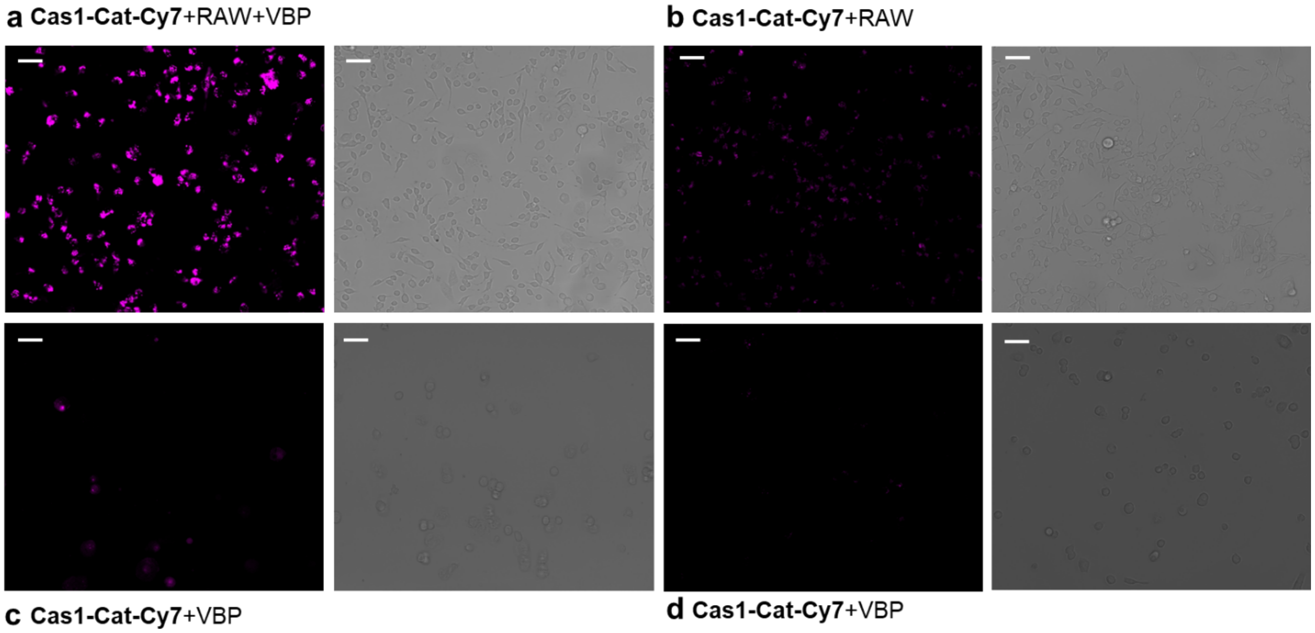


**Figure S1**. Fluorescence microscopy and brightfield images of EA.hy926 cells cocultured with RAW 264.7 cells labelled with **Cas1-Cat-Cy7.** Incubation protocols are described in Figure 3.

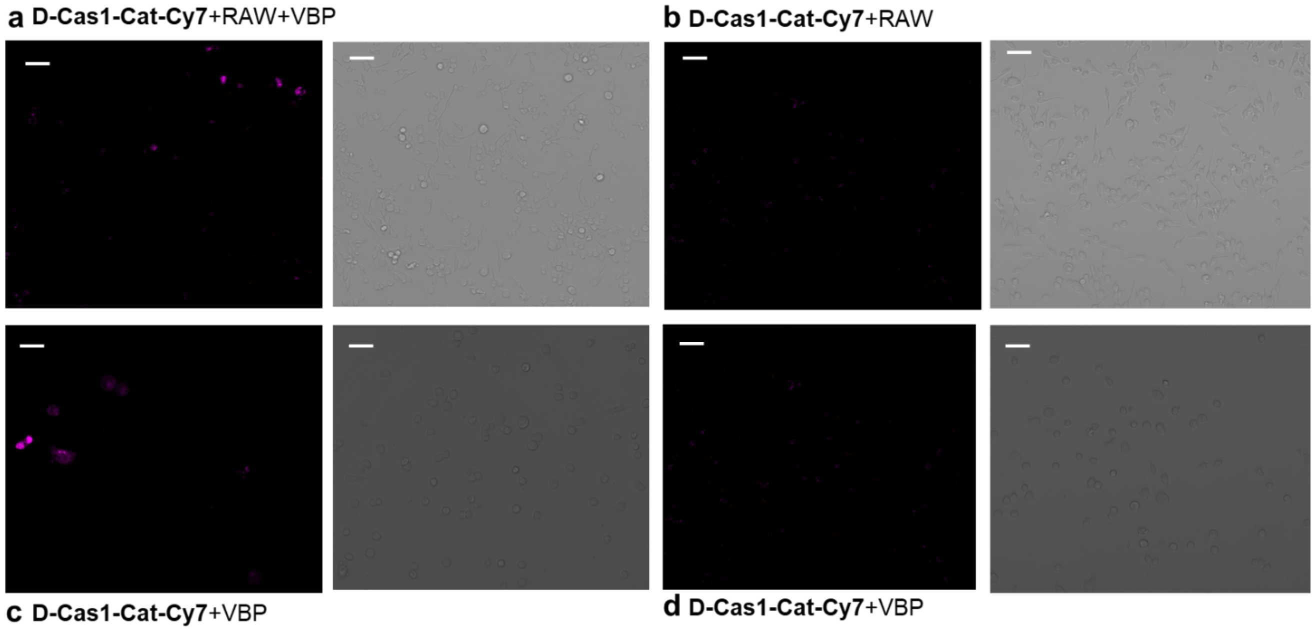


**Figure S2**. Fluorescence microscopy and brightfield images of EA.hy926 cells cocultured with RAW 264.7 cells labelled with **D-Cas1-Cat-Cy7.** Incubation protocols are described in Figure 3.

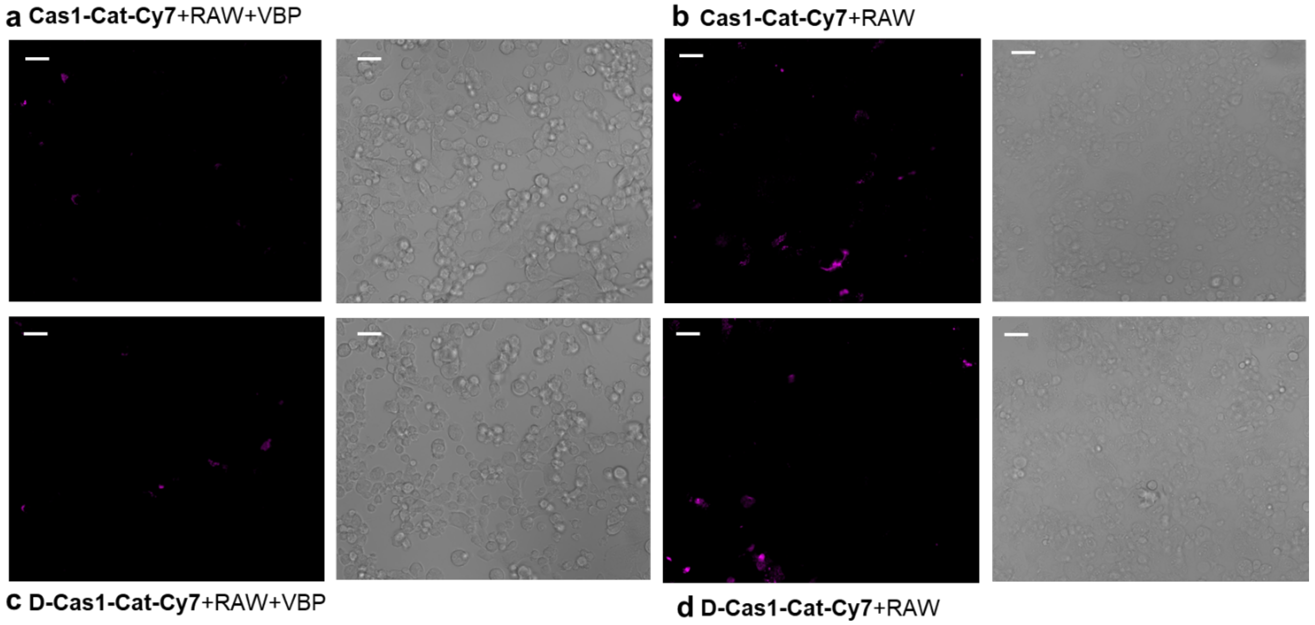


**Figure S3**. Fluorescence microscopy and brightfield images of EA.hy926 caspase1 KO cells cocultured with RAW 264.7 cells labelled with either **Cas1-Cat-Cy7** or **D-Cas1-Cat-Cy7.** a). Coculture of EA.hy926 KO and RAW 264.7 cells were incubated with Val-boroPro (50 μM) for 2h and then **Cas1-Cat-Cy7** (1 μM) was added and incubated for another 2h. b). Coculture of EA.hy926 KO and RAW 264.7 cells were incubated with **Cas1-Cat-Cy7** (1 μM) for 2h. c). Coculture of EA.hy926 KO and RAW 264.7 cells were incubated with Val-boroPro (50 μM) for 2h and then **D-Cas1-Cat-Cy7** (1 μM) was added and incubated for another 2h. d). Coculture of EA.hy926 KO and RAW 264.7 cells were incubated with **D-Cas1-Cat-Cy7** (1 μM) for 2h.

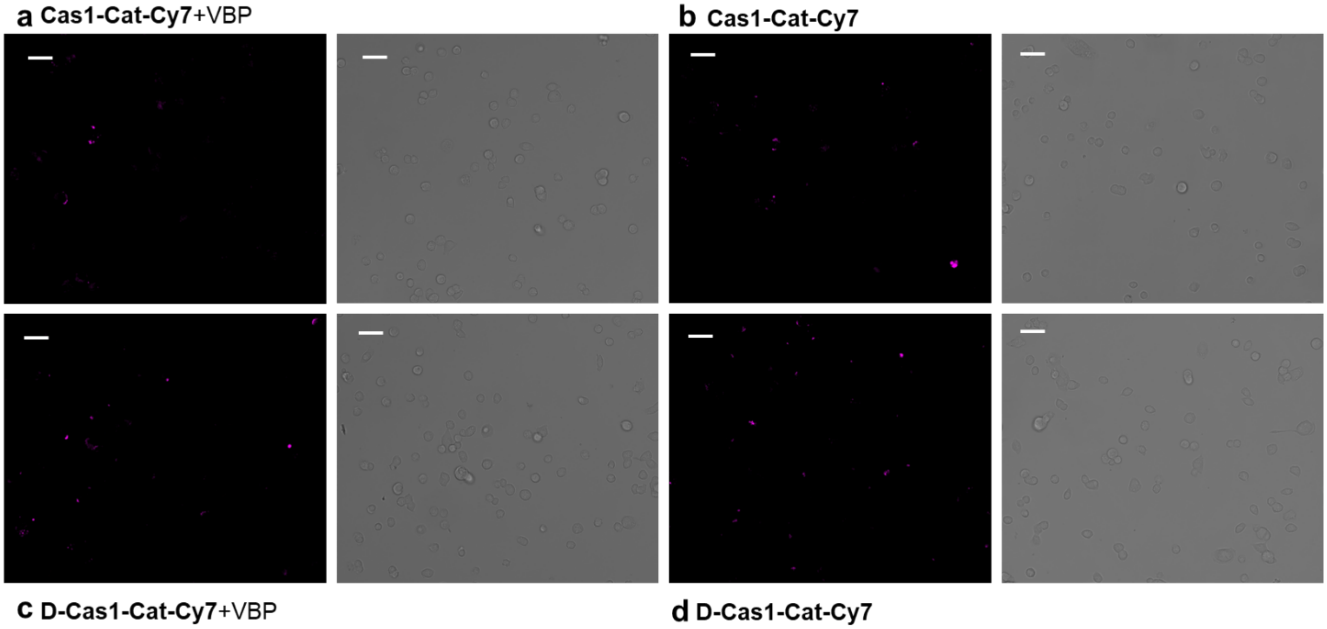


**Figure S4**. Fluorescence microscopy and brightfield images of EA.hy926 caspase1 KO cells labelled with either **Cas1-Cat-Cy7** or **D-Cas1-Cat-Cy7.** a). EA.hy926 KO cells were incubated with Val-boroPro (50 µM) for 2h and then **Cas1-Cat-Cy7** (1 µM) was added and incubated for another 2h. b). EA.hy926 KO cells were incubated with **Cas1-Cat-Cy7** (1 µM) for 2h. c). EA.hy926 KO cells were incubated with Val-boroPro (50 µM) for 2h and then **D-Cas1-Cat-Cy7** (1 µM) was added and incubated for another 2h. d). EA.hy926 KO cells were incubated with **D-Cas1-Cat-Cy7** (1 µM) for 2h.

### 5. LCMS of compounds

Intermediates containing Asp-Pro-tertLeu-Tyr showed two peaks in LCMS due to restricted rotation of the rigid structure.[3]

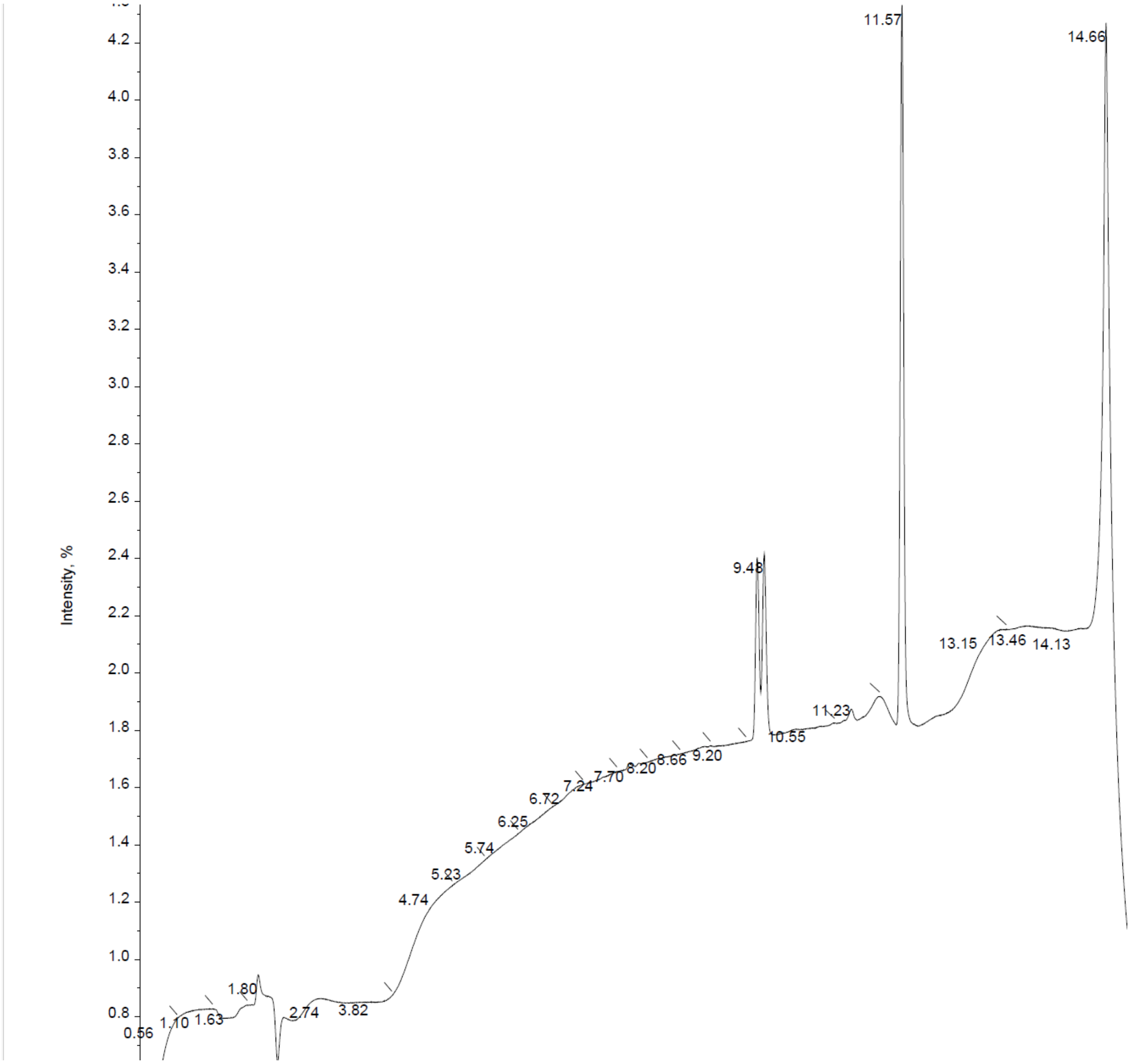


LC trace of intermediate **3**

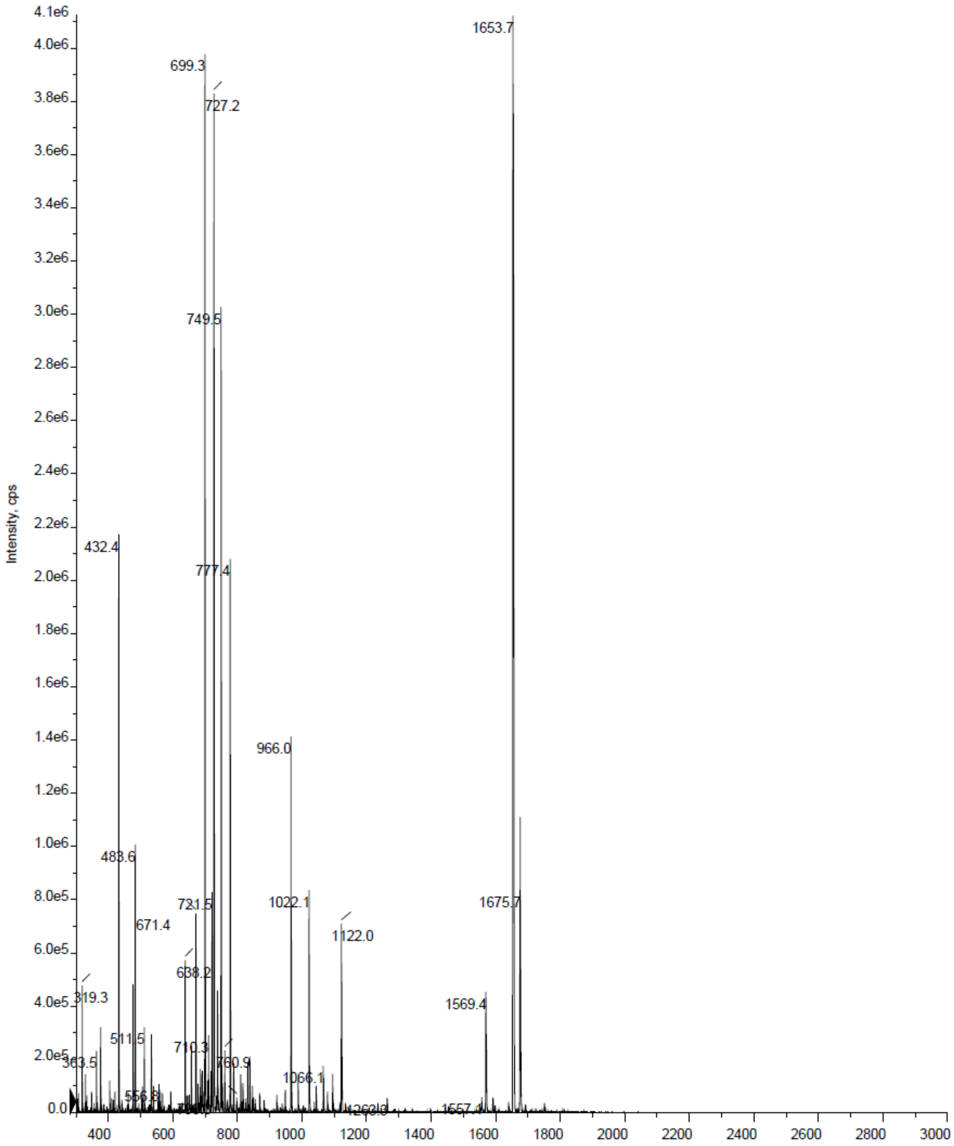


Mass of intermediate **3**

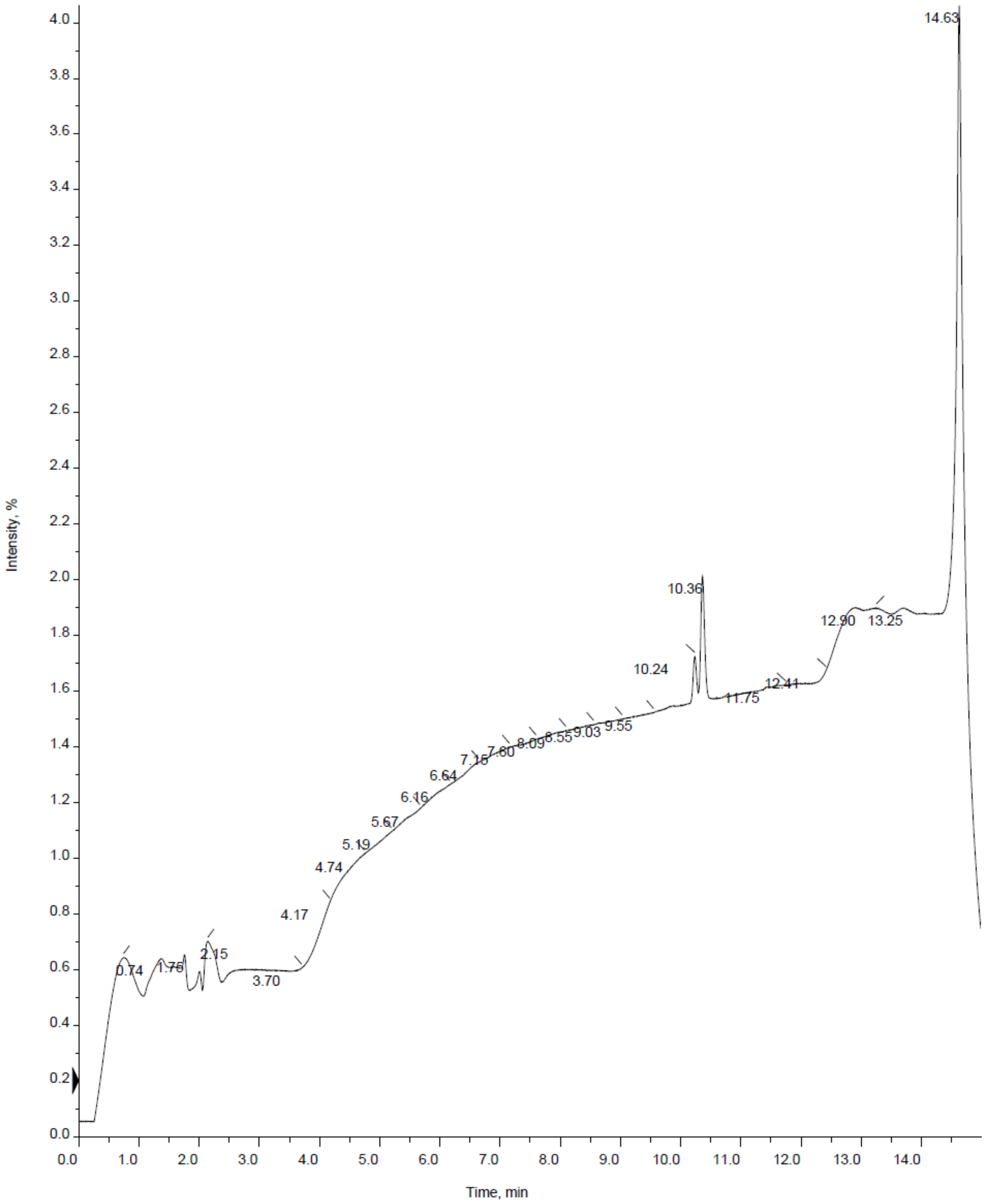


LC trace of intermediate **4**

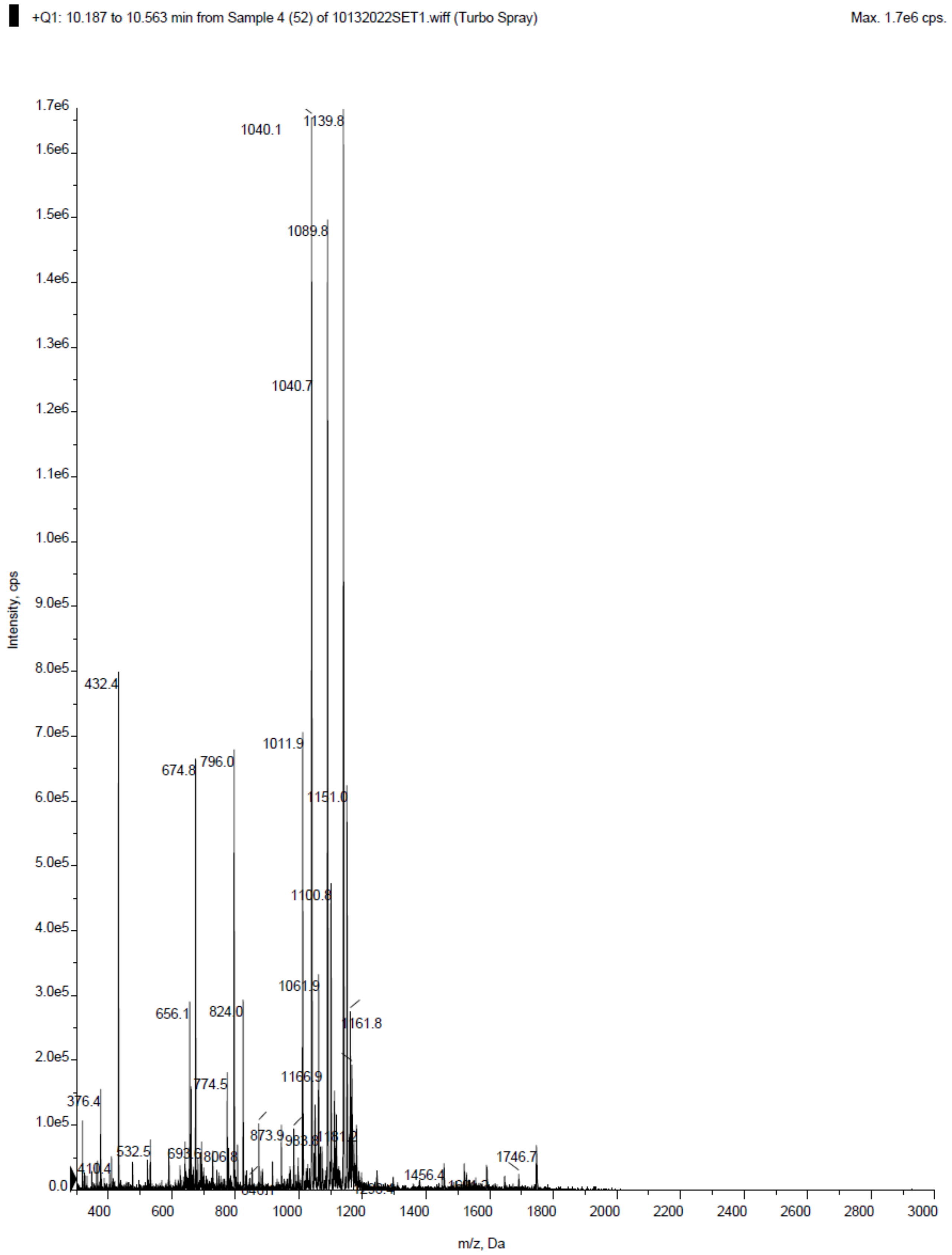


Mass of intermediate **4**

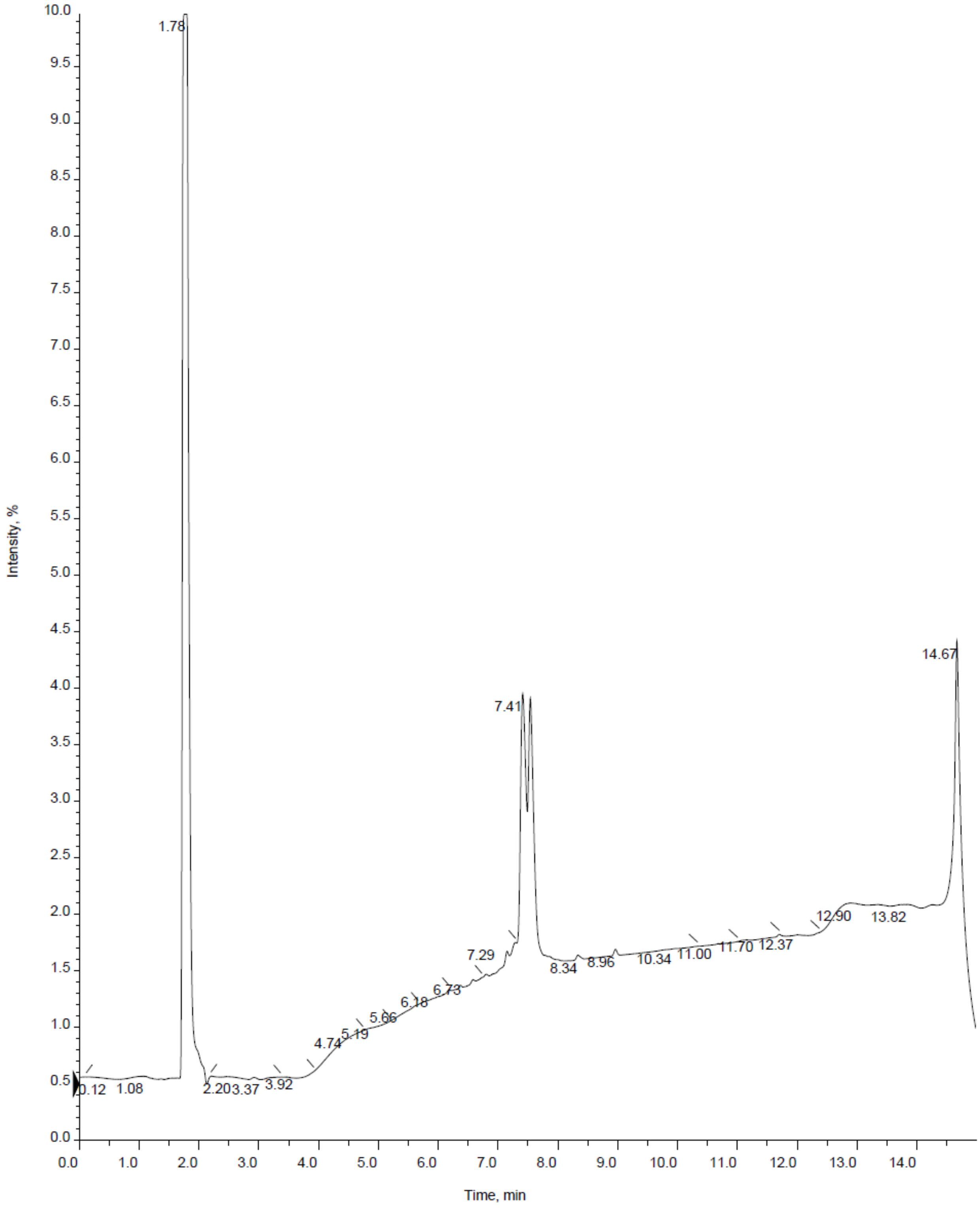


LC trace of **5** (**Cas1-Cat-Cy7**)

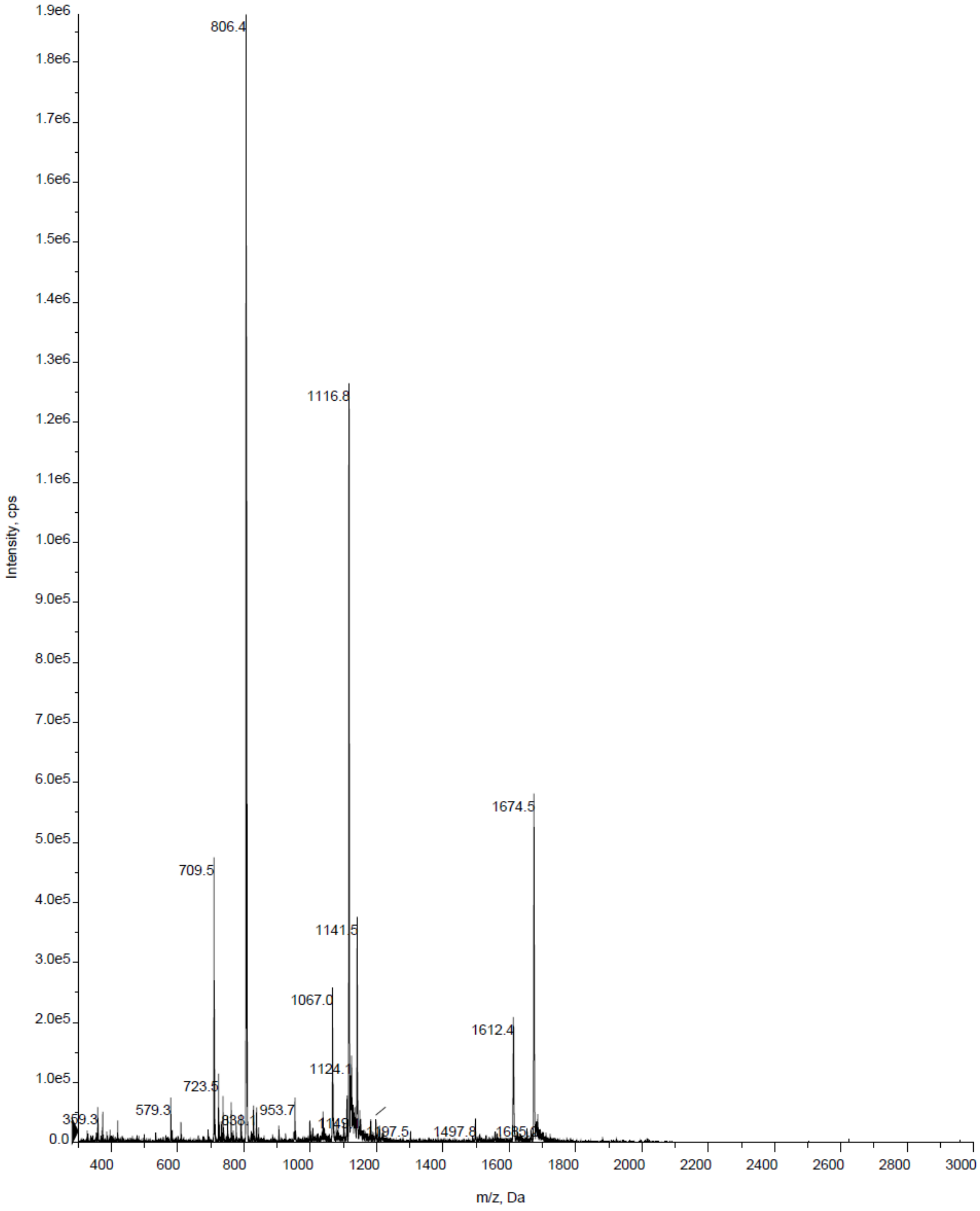


Mass of **5** (**Cas1-Cat-Cy7**)

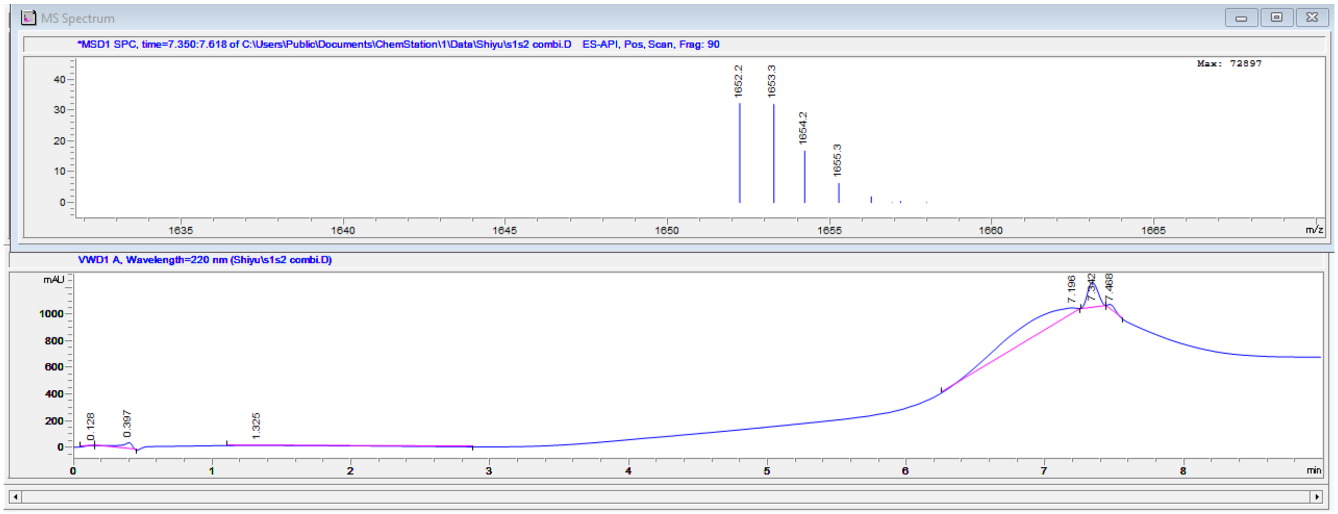


LCMS of **D-3**

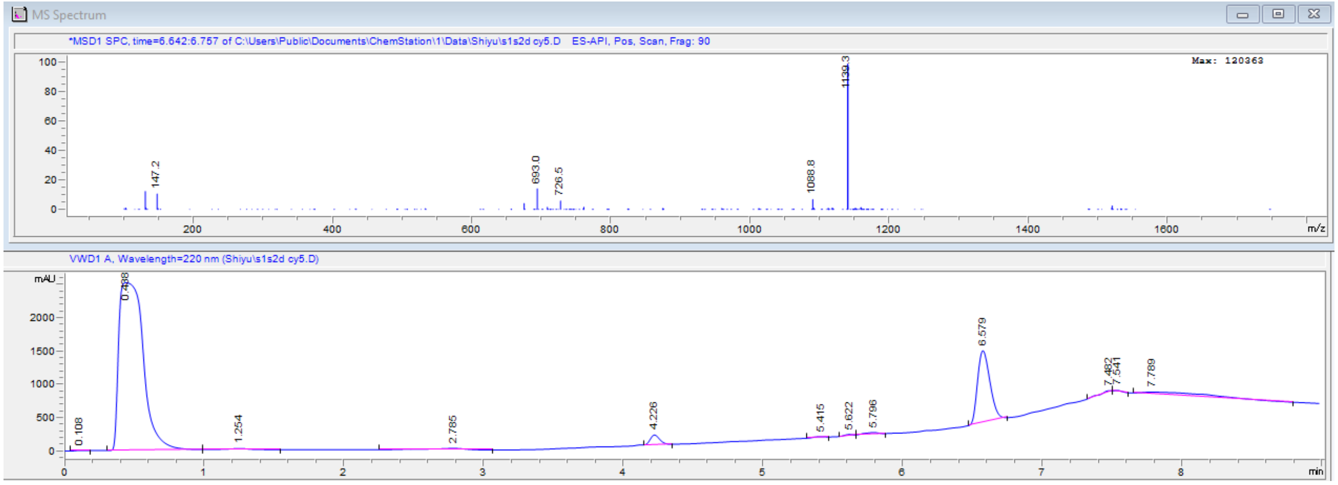


LCMS of **D-4**

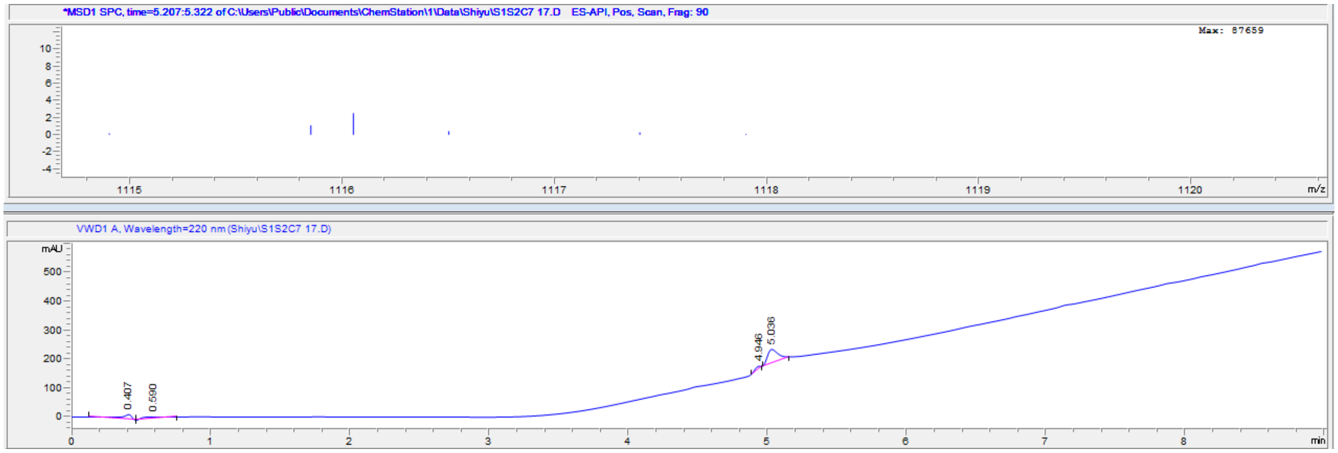


LCMS of **D-5** (**D-Cas1-Cat-Cy7**)

### 6. References

1. J. C. Widen, M. Tholen, J. J. Yin, A. Antaris, K. M. Casey, S. Rogalla, A. Klaassen, J. Sorger, M. Bogyo. AND-gate contrast agents for enhanced fluorescence-guided surgery. *Nat. Biomed. Eng.* **2021**, *5*, 264-277.
2. F. F. Faucher, K. J. Liu, E. D. Cosco, J. C. Widen, J. Sorger, M. Guerra, M. Bogyo. Protease activated probes for real-time ratiometric imaging of solid tumors. *Acs. Central. Sci.* **2023**, *9*, 1059-1069.
3. A. W. Puri, P. Broz, A. Shen, D. M. Monack, M. Bogyo. Caspase-1 activity is required to bypass macrophage apoptosis upon Salmonella infection. *Nat. Chem. Biol.* **2012**, *8*, 745-747.
